# A self-inactivating invertebrate opsin with resistance to retinal depletion optically drives biased signaling toward Gβγ-dependent ion channel modulation

**DOI:** 10.1101/2023.01.05.522954

**Authors:** Hisao Tsukamoto, Yoshihiro Kubo

## Abstract

Animal opsins, light-sensitive G protein-coupled receptors (GPCRs), have been utilized for optogenetic tools to control G protein-dependent signaling pathways. Upon G protein activation, the Ga and Gβγ subunits drive different intracellular signaling pathways, leading to complex cellular responses. For some purposes, Ga-, Gβγ-dependent signaling needs to be separately modulated, but these responses are simultaneously evoked due to the 1:1 stoichiometry of Ga and Gβγ. Nevertheless, we show temporal activation of G protein using a self-inactivating invertebrate opsin, *Platynereis* c-opsin1, drives biased signaling for Gβγ-dependent GIRK channel activation in a light-dependent manner by utilizing the kinetic difference between Gβγ-dependent and Ga-dependent responses. The opsin-induced transient Gi/o activation preferably causes activation of the kinetically-fast Gβγ-dependent GIRK channels rather than slower Gi/oα-dependent adenylyl cyclase inhibition. Although similar Gβγ-biased signaling properties were observed in a self-inactivating vertebrate visual pigment, *Platynereis* c-opsin1 needs fewer retinal molecules to evoke cellular responses. Furthermore, the Gβγ-biased signaling properties of *Platynereis* c-opsinl are enhanced by genetically fused with RGS8 protein which accelerates G protein inactivation. The self-inactivating invertebrate opsin and its RGS8-fusion protein can function as optical control tools biased for Gβγ-dependent ion channel modulation.

## Introduction

Optical control tools are necessary for optogenetic studies, to regulate specific cellular responses in a light-dependent manner^1^. Besides channelrhodopsins, animal opsins, light-sensitive G protein-coupled receptors (GPCRs), have been used as major optical control tools^2–4^ Animal opsins drive G protein- and arrestin-dependent signaling pathways upon illumination. In particular, Gi/o-coupled opsins are utilized as inhibitory tools via light-dependent Gi/oα-dependent inhibition of adenylyl cyclase and/or Gβγ-dependent modulation of G protein-activated inwardly rectifying potassium (GIRK) channels (see Fig. 1A) as well as voltage-gated calcium channels^5–8^. As optical control tools, animal opsins show much higher photosensitivity than channelrhodopsins, probably due to signal amplification via signaling cascades^6,9,10^.

**Fig. 1.**
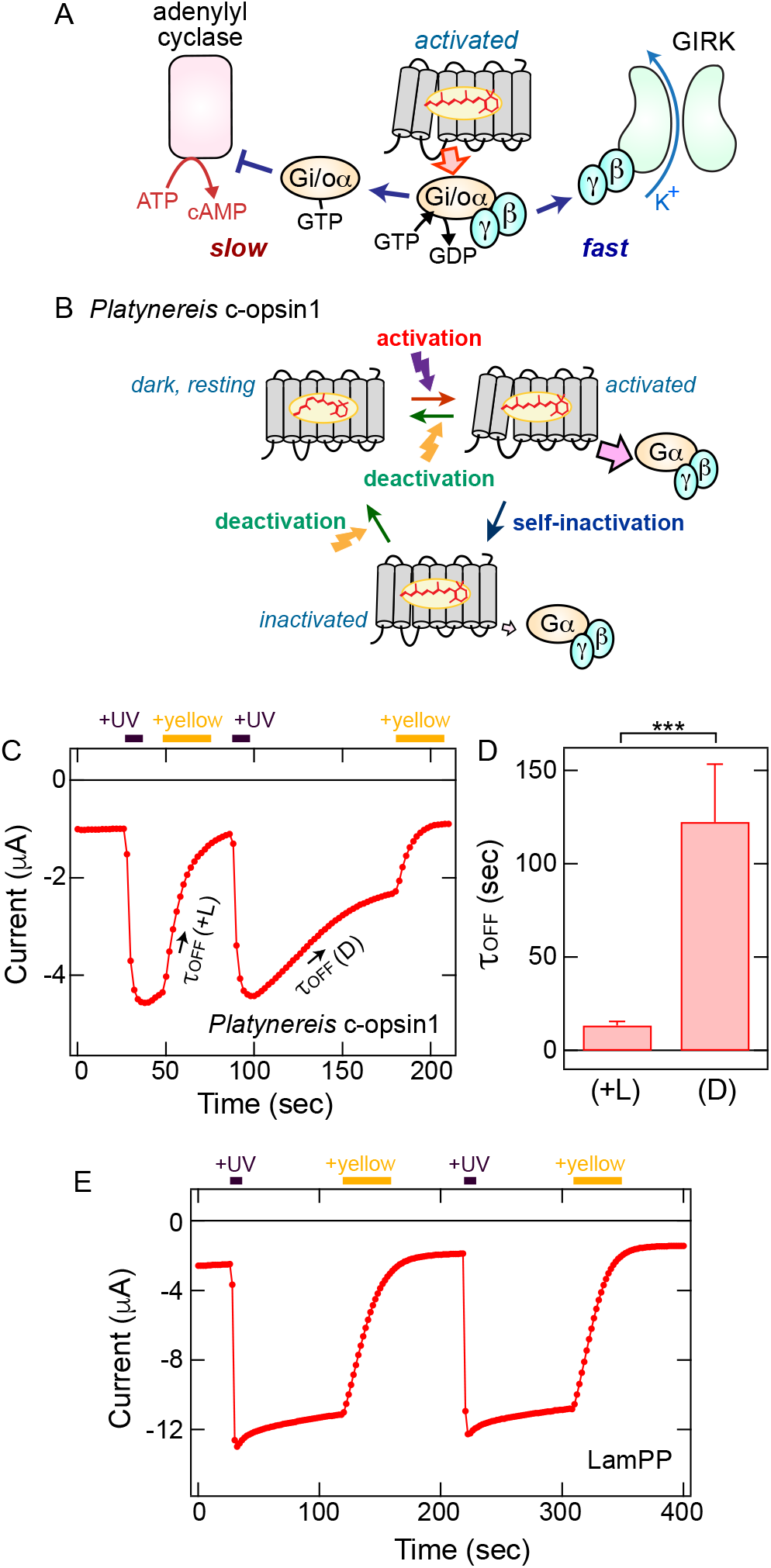
GIRK activation, inactivation and deactivation by c-opsin1 and LamPP. *A*, A schematic drawing to explain that an activated opsin molecule drives Gi/oα-dependent inhibition of adenylyl cyclase (slow response) and Gβγ-dependent activation of GIRK channel (fast response). *B*, A schematic drawing of activation/inactivation/deactivation of c-opsin1. *C*, UV-induced activation, visible (yellow) light-induced deactivation, and time-dependent (lightindependent) inactivation of GIRK1/GIRK2 channels by c-opsin1 in a *Xenopus* oocyte. Time lapse change of the current amplitude at −100 mV is plotted. Light stimulations were applied at the times indicated by the violet (UV light, 395-nm) and yellow (>500-nm light) bars. *D*, Comparison of the time constant (τ_off_) values of visible light-induced deactivation (+*L*) and timedependent inactivation (*D*) of the GIRK channels by c-opsin. Error bars indicate the S.D. values (n = 7). The mean ± S.D. values are 13.4±2.2 (+*L*) and 122.5±31.0 (*D*) seconds, respectively. The τ_off (+L)_ and τ_off (D)_ values are significantly different (\*\*\**P* = 7.8 × 10^-7^, t = −9.2957, d.f. = 12 by Student’s unpaired *t*-test). *E*, UV-induced activation, visible (yellow) light-induced deactivation of GIRK channels by LamPP. Time lapse change of the current amplitude at −100 mV is plotted. Light stimulations were applied at the times indicated by the violet (UV light, 395-nm) and yellow (>500-nm light) bars.

Animal opsins also possess some drawbacks as optical control tools. Most animal opsin-based optogenetic tools such as opto-XRs are developed using vertebrate visual pigments such as rhodopsins (rod opsins) or cone opsins^2–6^. Vertebrate visual pigments selectively bind *cis*-retinal isomers that are scarce outside the eyes and they are unidirectionally activated by light via *cis-* to *trans-* retinal isomerization (see supplemental Fig. S1A)^11,12^. In other words, vertebrate visual pigments require continuous supply of *cis*-retinal molecules as in visual photoreceptor cells. Recent optogenetic studies have proved that bistable opsins can overcome this issue^8,13–15^. Bistable opsins can bind *trans*-retinal isomer as well as *cis* isomers and are bidirectionally photo-interconvertible between stable (*cis*-retinal bound) resting/dark and (*trans*-retinal bound) activated forms (see Supplemental Fig. S1B)^3,16^. Another disadvantage of animal opsin as optical control tool stems from characteristics of GPCR signaling pathways. Activated GPCR promotes dissociation of G protein a and βγ subunits, each of which drives downstream pathways. Thus, animal opsin-induced cellular responses tend to be complicated^17^, but Ga- and Gβγ-dependent cellular responses do not seem to be separable due to the 1:1 stoichiometry of Ga and Gβγ subunits.

In this study, we aim to selectively drive Gβγ-dependent ion channel modulation by opsins that transiently activate G proteins (Fig. 1B). For this purpose, we utilize the kinetic difference between major Ga-and Gβγ-dependent cellular responses. In particular, Gβγ-mediated GIRK channel activation via the “membrane-delimited” process is much faster than Ga-mediated modulation of adenylyl cyclase regulating intracellular cAMP levels (Fig. 1A)^18–20^. This raises a possibility that transient Gi/o activation can evoke significant and temporal GIRK activation but not intracellular cAMP changes due to insufficient temporal integration of G protein activity upon receptor activation. We previously reported that an invertebrate Gi/o-coupled opsin (*Platynereis* c-opsin1)^21^ can be activated and deactivated by different color of light like typical bistable opsins, but the activated opsin is also spontaneously inactivated in a time-dependent manner^22^ (see Fig. 1C). This self-inactivating property (Fig. 1B) can be suitable for the temporal activation of Gi/o. Based on these characteristics, we hypothesize that the self-inactivating c-opsin1 preferably drives fast Gβγ-dependent GIRK channel response rather than slow Gi/oα-dependent cAMP reduction (Fig. 1A). In other words, we expect that c-opsin1 could function as a Gβγ-biased optical control tool.

Herein, we investigated light-dependent GIRK activation and adenylyl cyclase inhibition by c-opsin1 as well as other bistable and self-inactivating opsins. As expected, c-opsin1 and a vertebrate visual pigment having self-inactivating property predominantly drove Gβγ-dependent GIRK responses rather than Gi/oα-dependent cAMP reduction, whereas a typical bistable opsin produced robust Gβγ- and Ga-dependent responses. Also, c-opsin1 required less retinal molecules to function in comparison with the vertebrate visual pigment. The Gβγ-biased property of *Platynereis* c-opsin1 was enhanced by genetically combining with RGS8 to accelerate Gi/o inactivation. Taken together, we propose that the self-inactivating c-opsin1 and its fusion protein with RGS8 are suitable as optical control tools biased for kinetically-fast Gβγ-dependent signaling even under retinal depleted conditions.

## Results

We previously reported that *Platynereis* c-opsin1 can activate GIRK channels by UV light-dependent Gi/o activation^22^. The c-opsin1-induced GIRK current gradually decreased by termination of the illumination as well as irradiation with visible light, indicating that the G protein-activating state of c-opsin1 is transiently formed upon UV absorption. The activated form is inactivated in a time-dependent (light-independent) manner, and deactivated by visible light illumination (see Fig. 1B, C).

We expected that the transient Gi/o activation predominantly drives kinetically fast Gβγ-dependent GIRK activation rather than slow Gi/oα-dependent cAMP reduction response (Fig. 1A, see Introduction). To examine this, we quantified Gi/oα- and Gβγ-dependent cellular responses by c-opsin1 as well as other UV-sensitive opsins. We assessed Gi/oα-mediated adenylyl cyclase inhibition by Glosensor assay^23^ in mammalian cultured cells (COS-1 cells), and Gβγ-mediated GIRK activation by electrophysiological measurement in *Xenopus* oocytes^24,25^ (see Materials and Methods). The Ga- and Gβγ-dependent responses were compared between c-opsin1 and other UV-sensitive opsins, lamprey parapinopsin (LamPP) and mouse short-wavelength sensitive opsin (SWO), that are commonly used as optogenetic tools^6,9,13,15^.

### Gβγ-dependent GIRK channel responses by c-opsin1 and LamPP

In *Xenopus* oocytes, GIRK current increased by c-opsin1 upon UV illumination, and the increased GIRK current decayed in visible (yellow) light-dependent and time-dependent (light-independent) manners (Fig. 1C). Based on these data, we quantified τ_off (+L)_ and τ_off_ (D) values, which represent decay kinetics of the GIRK currents in light-dependent and independent manners, respectively (Fig. 1D). The respective τ_off_ values are 13.4±2.2 sec (by visible light-induced deactivation) and 122.5±31.0 sec (by self-inactivation) (Fig. 1D). We previously reported that the photoproduct of c-opsin1 is spectroscopically stable at least for 25 min after UV-absorption, indicating that the opsin holds the chromophore retinal^22^. Thus, the spontaneous inactivation of c-opsin1 is not caused by (all-*trans*) retinal release that occurs in vertebrate visual pigments (see Supplemental Fig. S1A). The visible light-induced deactivation was reported to be caused by photoisomerization of the retinal chromophore from *all-trans* to 11-*cis* configurations^22^ (Fig. 1B). In contrast to c-opsin1, a typical UV-sensitive bistable Gi/o-coupled opsin LamPP^26^ provoked sustained GIRK activation upon UV-illumination, and the GIRK current was turned off only by the visible light illumination (Fig. 1E), indicating that LamPP forms a stable G protein-activating state (see Supplemental Fig. S1B). The sustained GIRK activation by LamPP is consistent with previous optogenetic studies using LamPP^9,13,15,27^

### Gi/oα-dependent cAMP responses by c-opsin1 and LamPP

Glosensor assay of c-opsin1 and LamPP in COS-1 cells also demonstrates the transient nature of G protein-activating state of c-opsin1 (Fig. 2). Upon 10 sec UV illumination, LamPP induced larger decrease of intracellular cAMP levels^28^ than c-opsin1 (Fig. 2A). The tiny cAMP reduction by c-opsin1 is consistent with a previous study^29^. Subsequent 1, 2 and 4 min UV stimulations on c-opsin1 caused a transient cAMP reduction followed by a recovery in several minutes, and the same UV stimulations of LamPP caused additional transient cAMP decreases (Fig. 2A). The c-opsin1-induced transient cAMP responses can be explained by a temporal Gi/o activation (Fig. 1B). Due to the transient nature of the G protein-activating state, short time (10 sec) UV illumination of c-opsin1 caused a tiny cAMP reduction, and longer (several mins) continuous UV illuminations induced larger transient decreases of cAMP levels (Fig. 2A). In contrast, short time UV illumination of LamPP caused formation of the stable G protein-activating state to induce large and sustained cAMP reduction responses.

**Fig. 2.**
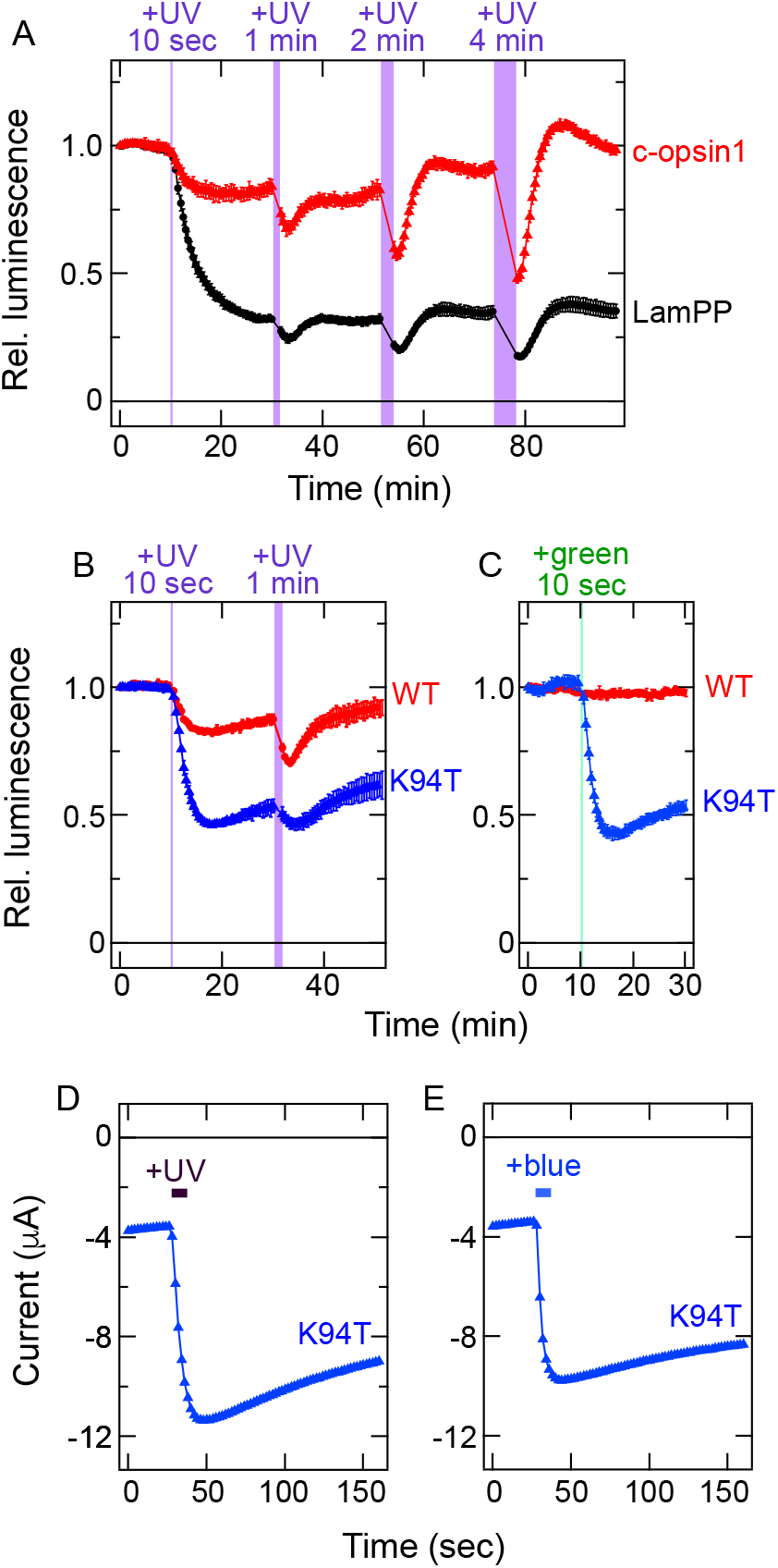
UV-dependent reduction of intracellular cAMP levels by c-opsin1. *A*, Light induced changes in cAMP biosensor (Glosensor) luminescence in COS-1 cells expressing *Platynereis* c-opsin1 (*red*) or LamPP (*black*). Luminescence levels are normalized to the value at the starting point (time = 0 min). Error bars indicate the S.D. values (n = 4). Violet bars show UV-light (375-nm) illumination and the duration of illumination are also indicated. Prior to the measurements, intracellular cAMP levels were elevated by application of 1 μM forskolin, an activator of adenylyl cyclase. *B* and *C*, Light induced changes in cAMP biosensor (Glosensor) luminescence in COS-1 cells expressing *Platynereis* c-opsin1 WT (*red*) or the K94T mutant (*blue*). Error bars indicate the S.D. values (n = 3). Violet and green bars show UV-light (375-nm) and green-light (500-nm) illuminations, respectively. As in panel *A*, 1 μM forskolin was added prior to the measurements. *D* and *E*, UV- or blue light-induced activation of GIRK1/GIRK2 channels by the K94T mutant of c-opsin1 in *Xenopus* oocytes. Time lapse changes of the current amplitude at −100 mV are plotted. (*blue*). Light stimulations were applied at the times indicated by the violet (UV light, 395-nm) blue (blue light, 470-nm) bars.

### K94T mutation-induced deceleration of the self-inactivation in c-opsin1

Interestingly, the K94T single mutant of c-opsin1, which causes a shift of absorption maximum from 383-nm to 491-nm^22^, showed sustained and larger responses upon UV or visible light illumination unlike WT (Fig. 2B, C), indicating that the mutation-induced spectral shift converts c-opsin1 so that it can be activated by visible light as well as UV light. The K94T mutant cannot be deactivated by light due to overlapping absorption spectra of resting and activated forms^22^. Interestingly, the K94T mutant evoked sustained GIRK current upon illumination (Fig. 2D, E). These results indicate that the limited cAMP responses by c-opsin1 is not due to efficient interaction with shut off proteins such as arrestin, because the position 94 is located at the retinal-binding site in the transmembrane domain and far from the G protein/arrestin interaction sites in the cytoplasmic domain.

### Transient dissociation of Gi trimer upon activation of c-opsin1

We directly assessed the transient Gi/o activation by c-opsin1 using the NanoBiT G protein dissociation assay^30,31^. In this assay, G protein subunits dissociation/association processes can be monitored based on a luciferase (NanoLuc) luminescence. Fig. 3A shows that after UV illumination of c-opsin1 WT, dissociated Gia and Gβγ subunits rapidly (re-) associated, but in the case of LamPP, these subunits remained dissociated. These results are consistent with our argument (Figs. 1, 2) that the c-opsin1 induces transient Gi/o activation upon UV illumination but LamPP continuously activates Gi/o. Due to the dead-time (about 10 sec) between illumination of the cells and luminescence measurement in our plate reader (see Materials and Methods), we could not detect the maximal dissociation level of the Gia and Gβγ upon activation of the c-opsin1 WT. Also, UV-illuminated c-opsin1 K94T mutant caused a longer dissociation of Gia and Gβγ than WT (Fig. 3A). These results are consistent with the differences in GIRK and cAMP responses induced by c-opsin WT and the K94T mutant (Fig. 2B-E).

**Fig. 3.**
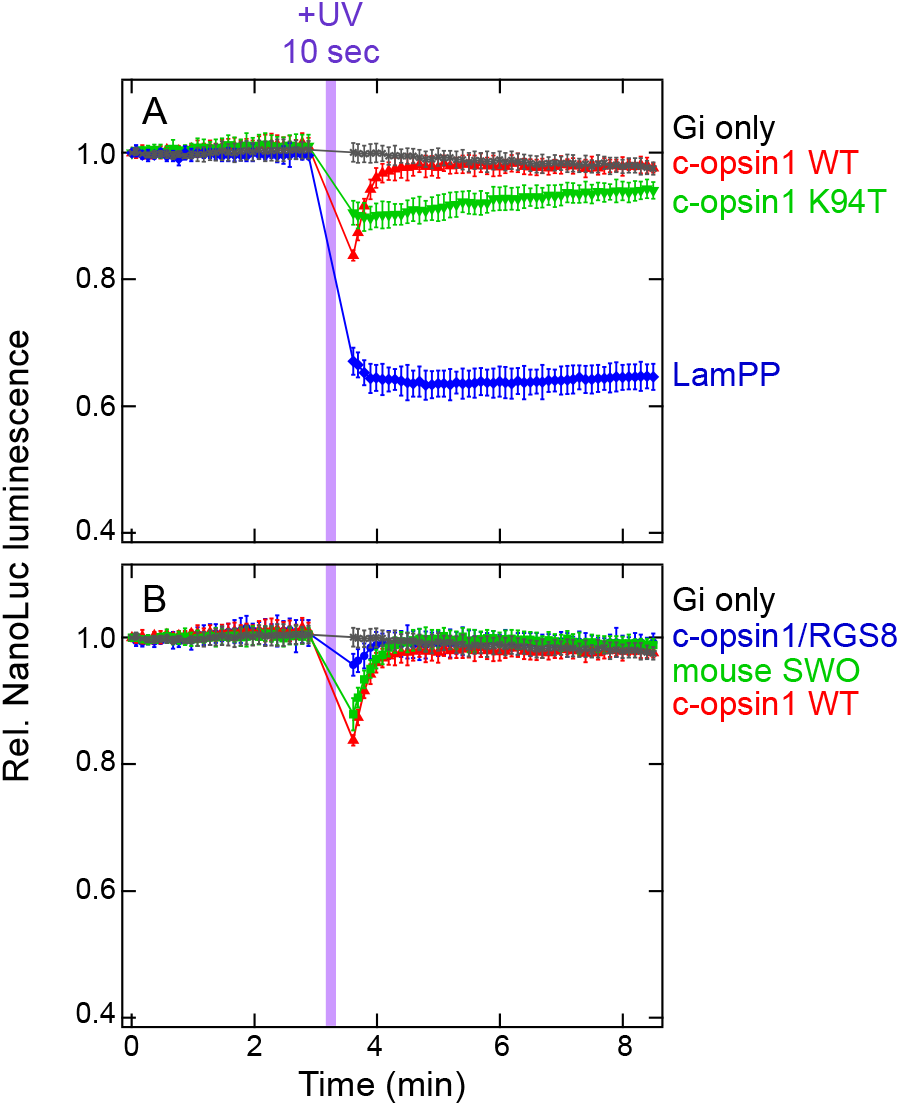
NanoBiT G protein dissociation assay on opsins used in this study. *A*, Kinetics of luminescence of NanoBiT Gi/o proteins upon UV-induced activation of c-opsin1 WT (*red*), the c-opsin1 K94T mutant (*green*), and LamPP (*blue*) in COS-1 cells. *B*, Kinetics of NanoLuc luminescence of NanoBiT Gi/o proteins upon UV-induced activation of c-opsin1 WT (*red*), the c-opsin1/RGS8 fusion protein (*blue*), and mouse SWO (*green*) in COS-1 cells. The same measurement was conducted without opsins (Gi only, *gray*). Data of “c-opsin1 WT” and “Gi only” are the same in panels *A* and *B*. Error bars indicate the S.D. values (n = 4). NanoLuc luminescence levels are normalized to the value at the starting point (time = 0 min). Violet bars indicate UV-light illumination and duration times of illumination are also indicated.

### GIRK and cAMP responses of mouse SWO with self-inactivating property

The cAMP responses of c-opsin1 are consistent with our hypothesis that temporal Gi/o activation preferably drives fast Gβγ-dependent GIRK activation rather than slow Gi/oα-dependent reduction of cAMP levels. Since photo-activated vertebrate visual pigments, in particular cone opsins, temporally activate G proteins and are spontaneously turned off by the retinal release^11,32^, we tested mouse SWO, a typical vertebrate UV-sensitive visual pigment^6^. In *Xenopus* oocytes, mouse SWO activated GIRK channels in a UV-dependent manner, and the activated GIRK current decayed in a time-dependent manner (τ_off (D)_ = 18.9±2.0 sec) (Fig. 4A, B). Also, Glosensor assay on the SWO showed that 10 sec irradiation with UV light caused a small transient cAMP reduction, and subsequent 1, 2, and 4 min UV stimulations caused larger transient responses followed by recovery (Fig. 4C) like the case of c-opsin1 (Fig. 2A). The NanoBiT G protein dissociation assay on mouse SWO also showed that mouse SWO, like *Platynereis* c-opsin1, induced transient Giαβγ dissociation upon UV illumination (Fig. 3B). Our data indicates that both c-opsin1 and mouse SWO predominantly drive fast GIRK activation via temporal activation of Gi/o, although the mechanisms underlying the self-inactivating nature are completely different between the two opsins. In other words, both *Platynereis* c-opsin1 and mouse SWO can act as optical control tools to preferably drive Gβγ-dependent ion channel modulation.

**Fig. 4.**
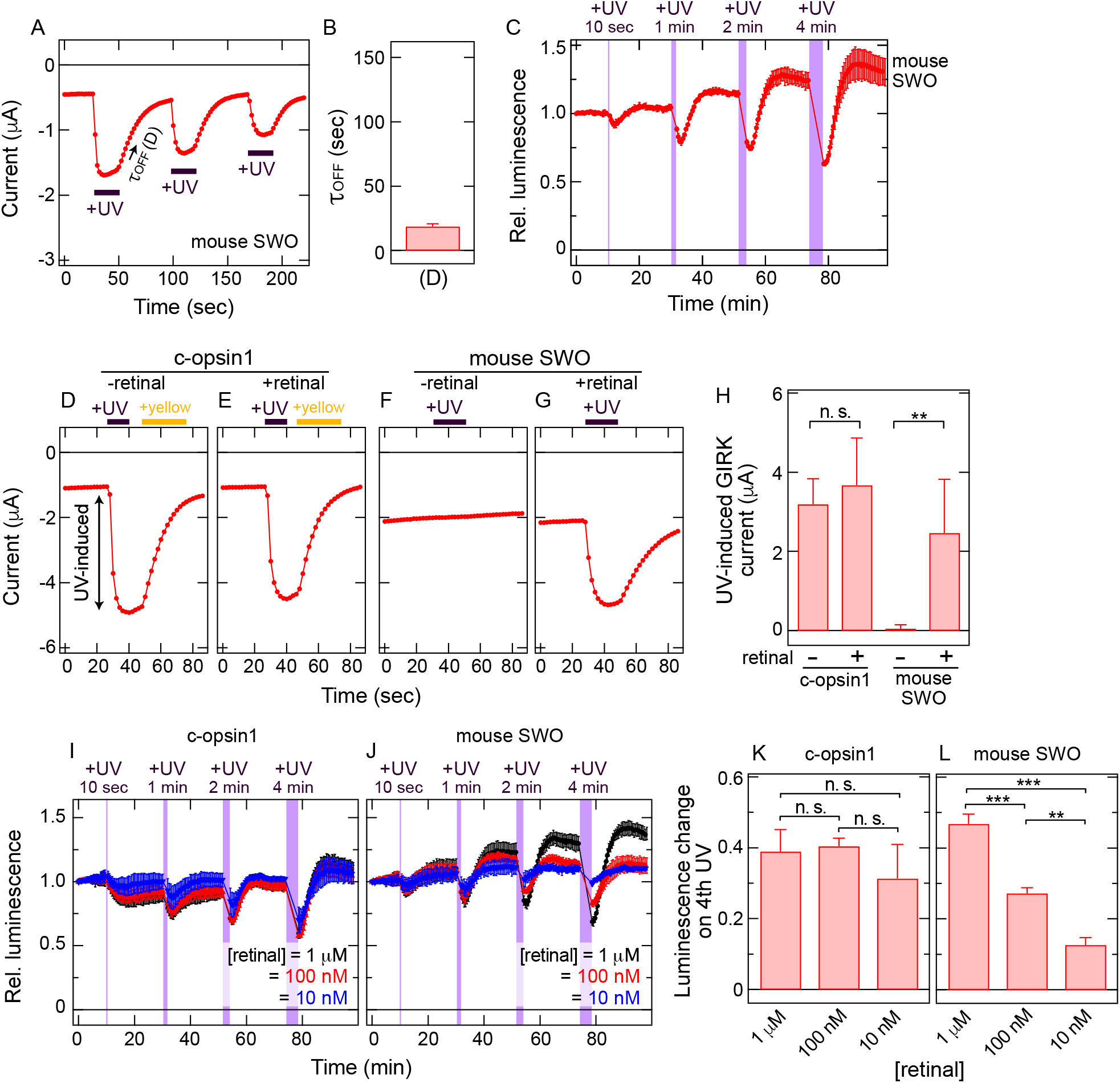
GIRK and cAMP responses driven by mouse SWO. *A*, UV-induced activation and time-dependent inactivation of GIRK1/GIRK2 channels by mouse SWO in a *Xenopus* oocyte. Time lapse change of the current amplitude at −100 mV is plotted. UV light (395-nm) stimulations were applied at the times indicated by the violet bars. *B*, Time constant (τ_off_) value of time-dependent inactivation of the GIRK channels by mouse SWO. Error bar indicates the S.D. value (n = 6). The mean ± S.D. value is 18.9±2.0 seconds. *C*, Light induced changes in cAMP biosensor (Glosensor) luminescence in COS-1 cells expressing mouse SWO. Luminescence levels are normalized to the value at the start point (time = 0 min). Violet bars indicate UV-light (395-nm) illumination and duration. Prior to the measurements, intracellular cAMP levels were elevated by application of 1 μM forskolin. *D – G*, UV-induced activation of GIRK1/GIRK2 channels by *Platynereis* c-opsin1 and mouse SWO in *Xenopus* oocytes in the presence and absence of exogenous retinal (1 μM). Time lapse changes of the current amplitude at −100 mV are shown. GIRK current amplitudes activated by *Platynereis* c-opsin1 with (*D*) or without (*E*) exogenous retinal, and those by mouse SWO with (*F*) or without (*G*) exogenous retinal, are shown. Light stimulations were applied at the times indicated by the violet (UV light, 395-nm) and yellow (>500-nm light) bars. The “UV-induced” current (*arrow*) indicates the amplitude of GIRK current increased by opsin activation. *H*, Comparison of UV-induced GIRK current upon activation of c-opsin1 and mouse SWO in the absence and presence of exogenous retinal molecules. Error bars indicate the S.D. values (n = 6, 7, 7, and 8, for c-opsin1-retinal, c-opsin1+retinal, mouse SWO-retinal, mouse SWO+retinal, respectively). The UV-induced GIRK current amplitudes are not significantly different for c-opsin1 (*n. s., P* = 0.26, t = −1.1822, d.f. = 11 by Student’s unpaired *t*-test) but significantly different for mouse SWO (\*\**P* = 0.00043, t = - 4.686, d.f. = 13 by Student’s unpaired *t*-test). *I* and *J*, Light induced changes in cAMP biosensor (Glosensor) luminescence in COS-1 cells expressing *Platynereis* c-opsin (*I*) or mouse SWO (*J*) in the presence of different concentrations of exogenous retinal. Luminescence levels are normalized to the value at the starting point (time = 0 min). *Black, red*, and *blue* data points indicate relative luminescence levels in the presence of 1 μM, 100 nM, and 10 nM exogenous 11-*cis*-retinal, respectively. Error bars indicate the S.D. values (n = 3). Violet bars indicate UV-light (375-nm) illumination and duration. Before the measurements, intracellular cAMP levels were elevated by application of 1 μM forskolin. *K* and *L*, Comparison of the luminescence change levels just after (*L*_after_) and before (*L*_before_) 4th (4 min) UV light (375-nm) illumination on COS-1 cells expressing *Platynereis* c-opsin1 (*K*) or mouse SWO (*L*). The luminescence change levels were calculated as [1 - *L*_after_/*L*_before_]. Error bars indicate the S.D. values (n = 3). In the presence of different concentrations of exogenous retinal, the luminescence changes of the 4th UV illumination are not significantly different for c-opsin1 (panel *K*, *n. s., P* = ***8.6 × 10^-5^, ***3.1 × 10^-6^, and **0.00047 for “1 μM” vs “100 nM”, “1 μM” vs “10 nM”, and “100 nM” vs “10 nM” respectively. F = 1.6, d.f. = 2, 6) but significantly different for mouse SWO (panel *L*, *n. s., P* = 0.96, 0.41, and 0.29 for “1 μM” vs “100 nM”, “1 μM” vs “10 nM”, and “100 nM” vs “10 nM” respectively. F = 182, d.f. = 2, 6). The statistical differences were evaluated using Tukey’s test following one-way ANOVA.

### Functionality of c-opsin in retinal depleted environments

*Platynereis* c-opsin1 and mouse SWO transiently activate Gi/o proteins as described above. In contrast, c-opsin1 can directly bind *cis*- and *trans*-retinal isomers, unlike vertebrate visual pigments including mouse SWO, and these retinal isomer-bound states are photo-interconvertible^22^ (Fig. 1B). The non-selective retinal-binding ability and photochemistry would make c-opsin1 resistant to retinal depletion (see Introduction). To test this, we measured the light-induced cellular responses by c-opsin1 and mouse SWO under retinal depleted conditions.

We quantified UV-induced GIRK current by c-opsin1 and mouse SW opsin in the absence or presence of exogenous 11-*cis*-retinal (1 μM) in *Xenopus* oocytes. Without exogenous retinal, both opsins can use only endogenous retinoids in oocytes. *Platynereis* c-opsin1 evoked UV-dependent GIRK current at similar levels regardless of addition of exogenous retinal (Fig. 4 D, E, H). In contrast, mouse SWO evoked little GIRK current upon UV illumination without exogenous retinal (Fig. 4F, G, H). These results clearly support our idea that c-opsin1 requires less exogenous retinal molecules to function. Also, we conducted Glosensor assay in COS-1 cells with different concentration of exogenous 11-*cis*-retinal (10 nM, 100 nM or 1 μM). A series of UV stimulations (10 sec, 1 min, 2 min and 4 min) were applied in this experiment, and UV-dependent decrease in cAMP levels were compared between c-opsin1 and mouse SWO. Consistent with electrophysiological data from oocytes, the Glosensor assay data showed that c-opsin1-induced cAMP reduction was less affected by the amount of exogenous retinal (Fig. 4I, K). Mouse SWO-induced cAMP responses dramatically decreased in retinal depleted conditions, and in particular, responses to the last 4 min UV illumination after 10 sec, 1 min, and 2 min UV irradiation were most affected in comparison with retinal-abundant conditions (Fig. 4 J, L). This is likely because mouse SWO, a typical vertebrate visual pigment, selectively binds *cis*-retinal isomers and releases isomerized (*trans-*) retinal after photoactivation (Supplemental Fig. S1A), and a series of activation by repeated and prolonged UV illumination consumes available *cis*-retinal molecules. In contrast, c-opsin1 holds both *cis*- and *trans*-retinal isomers, and upon UV illuminations both isomers are reversibly interconverted (Fig. 1B). Since c-opsin1 continuously uses the retinal molecule once bound to the protein, the opsin can evoke light-dependent responses even in a retinal depleted environment, unlike vertebrate visual pigment including mouse SWO (see Discussion).

### Enhancement of Gβγ-biased signaling properties of c-opsin1 by combination with RGS8

As c-opsin1 can preferably drive Gβγ-dependent GIRK activation (Figs. 1, 2), we concluded that the Gβγ-biased signaling property of c-opsin1 is caused by its self-inactivating property. However, the opsin also showed some residual UV-dependent cAMP reduction (Fig. 2A). In order to minimize the residual Gi/oα-dependent responses by c-opsin1, we tried to accelerate the shut-off of Gi/o signaling. RGS proteins are known to accelerate GTP hydrolysis in trimeric G proteins and termination of G protein-mediated signal transduction^33^. In particular, rat RGS8 was reported to efficiently accelerate both turn-off as well as turn-on kinetics of GIRK activation by Gi/o-coupled receptors^25^. Based on these insights, we engineered *Platynereis* c-opsin1 to be genetically fused with rat RGS8 on the C-terminus of the opsin (see Materials and Methods and Supplemental Data) to enhance temporal activation of G proteins (Fig. 5A). We expected that the fused protein would show enhanced Gβγ-biased signaling properties.

**Fig. 5.**
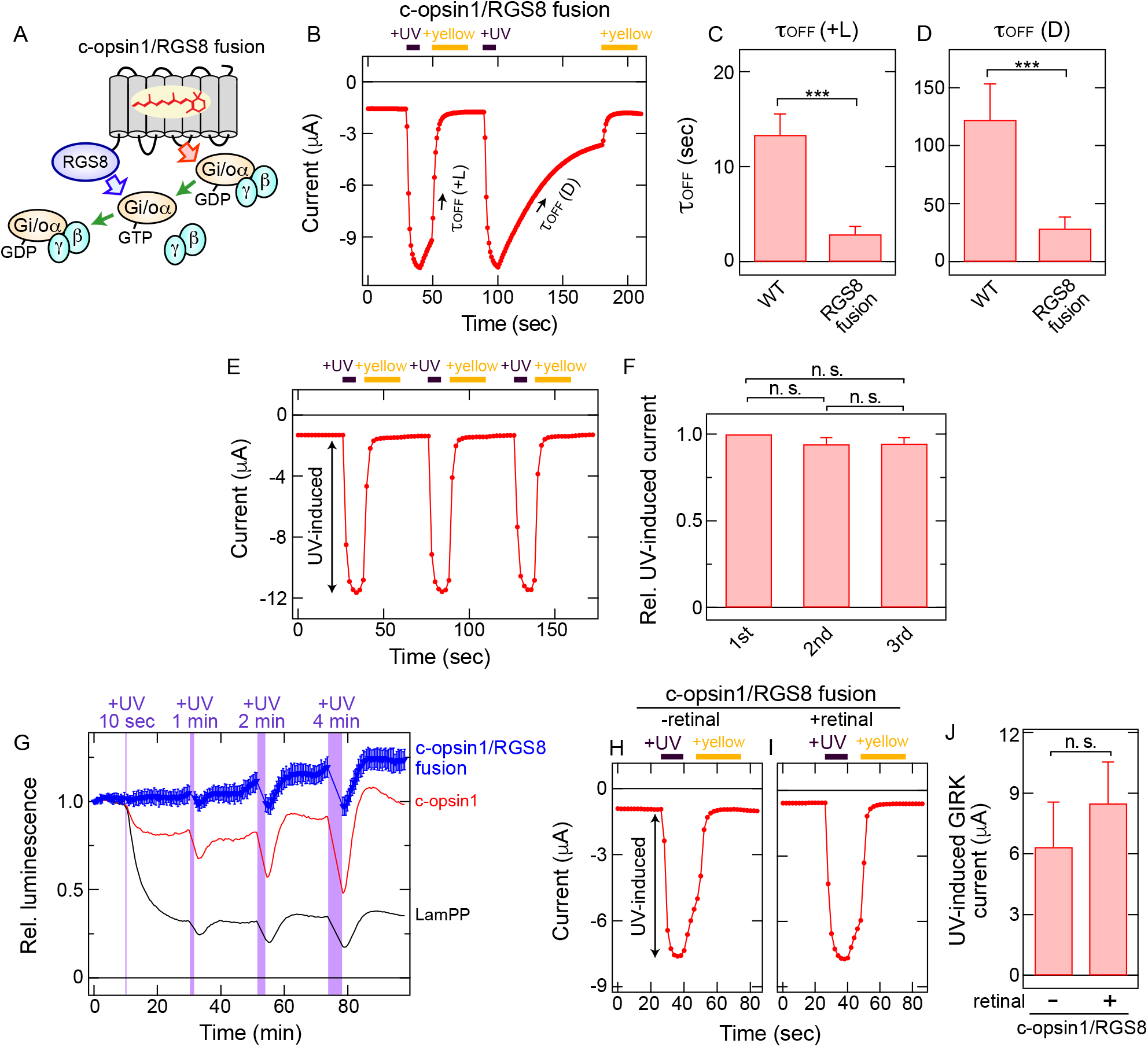
Light-induced GIRK and cAMP responses by c-opsin1/RGS fusion protein. *A*, A schematic drawing how the fusion protein modulates the activity of c-opsin1. The RGS8 moiety genetically fused on the C-terminus of c-opsin1 is expected to accelerate GTP hydrolysis in the Gi/oa that is activated by the c-opsin1 moiety. The acceleration by Gi/o inactivation would enhance Gβγ biased activity of c-opsin1. *B*, UV-induced activation, visible (yellow) light-induced deactivation, and time-dependent (light-independent) inactivation of GIRK1/GIRK2 channels by the c-opsin1/RGS8 fusion in a *Xenopus* oocyte. Time lapse change of the current amplitude at - 100 mV is shown. Light stimulations were applied at the times indicated by the violet (UV light) and yellow (>500-nm light) bars. In this measurement, data points are measured every 1 sec to analyze the fast kinetics (see Materials & Methods). *C* and *D*, Comparison of the time constant (τ_off_) values of visible light-induced deactivation (+*L*) (panel *C*) and time-dependent inactivation (*D*) (panel *D*) of the GIRK channels between c-opsin1 WT and the c-opsin1/RGS8 fusion protein. Error bars indicate the S.D. values (n = 7 for τ_off (+L)_ and n = 6 for τ_off (D)_ values of the RGS8 fusion). The mean ± S.D. values of the c-opsin1/RGS8 fusion protein are 2.9±0.8 (+*L*) and 28.8±9.9 (*D*) seconds, respectively. The τ_off (+L)_ and τ_off (D)_ values of c-opsin1 WT are the same as those in Fig. 1D. Both τ_off (+L)_ (\*\*\**P* = 5.6 × 10^-8^, t = 11.829, d.f. = 12 by Student’s unpaired *t*-test) and τ_off (D)_ (\*\*\**P* = 2.1 × 10^-5^, t = 7.0713, d.f. = 11 by Student’s unpaired *t*-test) values are significantly different between c-opsin1 WT and the c-opsin1/RGS8 fusion protein. *E*, Repeated GIRK activation and deactivation by the c-opsin/RGS8 fusion protein. *F*, Comparison of relative UV-induced GIRK current amplitudes upon repeated UV-induced activation and yellow light-induced deactivation. The labels “1st”, “2nd”, and “3rd” indicate the numbers of UV illumination. The amplitudes are normalized to the amplitudes upon the 1st UV illuminations. Error bars indicate the S.D. values (n = 4). The relative UV-induced GIRK current amplitude were not significantly different among the data sets (*n. s., P* = 0.052, 0.075, and 0.97 for “1st” vs “2nd”, “1st” vs “3rd”, and “2nd” vs “3rd” respectively. F = 4.7, d.f. = 2, 9). The statistical differences were evaluated using Tukey’s test following one-way ANOVA. *G*, UV light induced changes in cAMP biosensor (Glosensor) luminescence in COS-1 cells expressing the c-opsin/RGS8 fusion protein (*blue*). Error bars indicate the S.D. values (n = 4). The data of *Platynereis* c-opsin1 (*red*) and LamPP (*black*) are adopted from Fig. 2A. Luminescence levels are normalized to the value at the starting point (time = 0 min). Violet bars represent UV-light illumination and duration. Before the measurements, intracellular cAMP levels were elevated by application of 1 μM forskolin. *H* and *I*, UV-induced activation of GIRK1/GIRK2 channels by the c-opsin1/RGS8 fusion protein in *Xenopus* oocytes in the presence and absence of exogenous retinal (1 μM). Time lapse changes of the current traces at −100 mV are shown. GIRK current traces with (*H*) or without (*I*) exogenous retinal are shown. Light stimulations were applied at the times indicated by the violet (UV light, 395-nm) and yellow (>500-nm light) bars. The “UV-induced” current (*arrow*) indicates the amplitude of GIRK current increased by opsin activation. *J*, Comparison of UV-induced GIRK current amplitude upon activation of the c-opsin1/RGS8 fusion protein in the absence and presence of exogenous retinal molecules. Error bars indicate the S.D. values (n = 6). The UV-induced GIRK current amplitudes are not significantly different for the c-opsin1/RGS8 fusion (*n. s. P* = 0.11, t = −1.7598, d.f. = 10 by Student’s unpaired *t*-test).

The c-opsin1/RGS8 fusion protein expressed in *Xenopus* oocytes actually showed faster shut-off of GIRK activation and smaller τ_off (D)_ and τ_off (+L)_ values than those of c-opsin1 without RGS8 (Fig. 5B, C, D). By termination of UV illumination, the fusion protein spontaneously stopped the GIRK activation with τ_off (D)_ of 28.8±9.9 sec, and a visible light stimulation of the fusion protein quickly shut off the GIRK responses with τ_off (+L)_ of 2.9±0.8 sec (Fig. 5, C, D). These results suggest that the rate limiting step in the decay of GIRK activation by c-opsin1 is deactivation of G proteins via GTP hydrolysis. Notably, the τ_off (+L)_ value of the c-opsin1/RGS8 fusion protein was faster than the τ_off (D)_ value of mouse SWO, whereas vertebrate cone opsins such as the SWO showed rapid self-inactivation via retinal release^6,11,32^ (Fig. 5C and Supplemental Fig. S1A). In other words, the c-opsin1/RGS8 fusion protein can modulate GIRK channels more quickly than mouse SWO. The UV-induced activation and rapid visible light-induced deactivation of GIRK responses can be repeated, similarly to the function of intact c-opsin1 (Fig. 5E, F). The electrophysiological experiments proved that fusion with RGS8 enhances the characteristics of c-opsin1 to transiently and repeatedly activate Gi/o. The NanoBiT G protein dissociation assay on the c-opsin1/RGS8 fusion protein showed little UV-induced dissociation and the small signal rapidly diminished (Fig. 3B). We speculate that during the ~10 sec dead-time between the UV-illumination and luminescence measurement, the dissociated Gi trimer has reassociated more rapidly in the case of the c-opsin1/RGS fusion, than that of intact c-opsin1, leading to the observation of the very limited dissociation signal.

We next conducted Glosensor assay on the c-opsin1/RGS8 fusion protein to confirm that the fusion with RGS8 minimizes residual Ga-dependent responses. As expected, the c-opsin1/RGS8 fusion protein showed smaller residual cAMP reduction responses than the c-opsin1 WT (Fig. 5G). The fusion protein modulated GIRK channels in *Xenopus* oocyte in the absence of exogenous retinal like the case of the c-opsin1 WT (Fig. 4H), indicating that ligation with RGS8 does not interfere characteristic resistance of c-opsin1 to retinal depletion (Fig. 5H, I, J). Taken together, the c-opsin1/RGS8 fusion protein is our best optical control tool biased for Gβγ-dependent ion channel modulation with minimal residual activity of Gi/oα-dependent cAMP reduction responses (Supplemental Table).

## Discussion

In this study, we characterized *Platynereis* c-opsin1 as an optical control tool biased for Gβγ-dependent ion channel responses with small residual Ga-dependent enzymatic responses (Figs. 1, 2). Also, we showed c-opsin1 can function in retinal-depleted environments. Furthermore, we enhanced the Gβγ-biased property of c-opsin1 by fusion with RGS8 (Fig. 5). Here, we summarize characteristics of the c-opsin1-based optical control tools and their potential optogenetic applications.

### Self-inactivating property of Platynereis c-opsin1 correlating to Gβγ-biased signaling activity

*Platynereis* c-opsin1 transiently forms G protein-activating state(s) followed by spontaneous (light-independent) inactivation (Figs. 1C, 1D, 2A), although typical bistable opsins such as LamPP^26^ form stable G protein-activating states evoking sustained cellular responses (Figs. 1E, 2A). On the other hand, c-opsin1 can bind *cis*- and *trans*-retinal isomers directly like typical bistable opsins, and the *cis*- and *trans*-retinal bound states are spectroscopically stable and photo-interconvertible^22^. The light-induced deactivation via *trans* to *cis* photoisomerization of retinal is faster than spontaneous (light-independent) inactivation (Figs. 1C, 5B). Based on these data, we summarize a potential scheme describing photo-activation, spontaneous inactivation, and light-dependent deactivation of c-opsin1 in Fig. 1A. These processes are remarkably different from those of typical bistable opsins with a stable G protein-activating form (Supplemental Fig. S1B) and vertebrate visual pigments such as mouse SWO involving *trans*-retinal release and *cis-* retinal binding (Supplemental Fig. S1A). The light-independent self-inactivation process in c-opsin1 is rather similar to the sequential transition from metarhodopsin-II (fully active) to metarhodopsin-III (partially active) in vertebrate visual pigments^34^.

The self-inactivating nature of c-opsin is important to preferably drive fast Gβγ-dependent GIRK responses rather than slow Gi/oα-dependent cAMP responses. This is probably because the amplitude of fast GIRK response depends on maximal G protein activity levels upon light absorption, but that of slow cAMP response depends on integration of G protein activity on each time point after opsin activation. Maximal G protein activation levels are comparable between c-opsin1 and LamPP, but temporal integration of the G protein activation by the self-inactivating c-opsin1 is smaller than that by LamPP.

Light-independent self-inactivating property was also observed in another invertebrate Gi/o-coupled opsin *Anopheles* Opn3^35^. In contrast to *Platynereis* c-opsin1, *Anopheles* Opn3 induces significant cAMP reduction responses^35^, probably because its inactivation kinetics is much slower than that of *Platynereis* c-opsin1. Similarly, the K94T substitution in c-opsin1 slows the light-independent inactivation (Figs. 2D, 2E, 3A) leading to a larger light-dependent cAMP reduction than wild-type (Fig. 2, B, C). These insights suggest that amino acid substitutions in opsins to accelerate self-inactivating kinetics would enhance the Gβγ-biased signaling property.

### Functionality of Platynereis c-opsin1 in retinal depleted environment

As summarized in Figs. 4H, 4K, and 5J, *Platynereis* c-opsin1 and its fusion with RGS8 protein maintain their photosensitivity even under retinal depleted conditions. This is probably because c-opsin1 possesses non-selective binding ability for *cis-* and *trans-* retinal isomers and holds the retinal upon illumination, unlike vertebrate visual pigments such as mouse SWO^22^ (see Introduction). Notably, *Anopheles* Opn3 can directly bind 13-*cis*-retinal, which is easily generated from all-*trans*-retinal via thermal isomerization, and it can function under retinal depleted conditions, too^35^. Taken together, binding ability for *all-trans* and/or 13-*cis* retinal isomers and continued use of the once bound retinal molecules are preferable for opsin to function in retinal depleted environments.

Many tissues/organs contain very limited amounts of retinal molecules. The pineal organ in mammals expresses visual pigments, but are not photo-sensitive due to a lack of sufficient retinal supply^36,37^. Peripheral tissues such as the outer ears and vibrissal pads of mice cannot provide sufficient retinal molecules to endogenous opsin (Opn5) molecules *ex vivo*^38^. In insects such as *Drosophila*, limited amounts of retinal is available outside the eyes^39^. Also, in some optogenetic studies, exogeneous retinal molecules are added to make optogenetic tools functional^40,41^, although many neural cells in the mammalian brain contains sufficient retinal molecules for channelrhodopsins to function^1^. Some invertebrate opsins such as *Platynereis* c-opsin1 and

*Anopheles* Opn3 will expand the applicability of optogenetic techniques into various tissues/organs containing limited retinal molecules. Future studies will identify other invertebrate opsins that can function in retinal depleted tissues.

### Advantages of c-opsin1-based optical control tools

Gi/o-coupled bistable opsins are utilized as inhibitory optogenetic tools to evoke a sustained inhibitory GIRK current. Channelrhodopsin-based inhibitory tools need continuous illumination to generate sustained inhibitory current (but see ref. 42^42^). Thus, the self-inactivation in c-opsin1 seems to jeopardize the advantage, but here we proved that the property is rather beneficial for the selective control of Gβγ-dependent ion channel modulation. Vertebrate visual pigments including mouse SWO (Fig. 4) have been used as optogenetic tools to drive Gi/o signaling pathways^5,6^. Our data in Fig. 4 shows that mouse SWO can preferably drive Gβγ-dependent GIRK activation like c-opsin1, but the SWO needs a larger supply of retinal molecules than c-opsin1. C-opsin1-based optical control tools is suitable to drive Gβγ-dependent ion channel responses in retinal-depleted tissues (Supplemental Table).

Our data in Fig. 5, c-opsin1 genetically fused with RGS8 enhances the biased activity of c-opsin1 for Gβγ-dependent GIRK modulation by accelerating kinetics, maintaining the resistance for retinal depletion. We propose that the c-opsin1/RGS8 fusion protein is our best optical control tool biased for Gβγ-dependent ion channel modulation with a minimal residual Gi/oα-dependent activity (Supplemental Table). Fusion with RGS proteins could change signaling properties of other animal opsins, leading to development of valuable optical control tools.

### Potential of the c-opsin1-based Gβγ-biased optical control tools to expand optogenetics

Gi/o-coupled animal opsins are utilized as inhibitory optogenetic tools, and Gi-DREADD is used for an inhibitory chemogenetic tool^43^. They suppress neural activities via Gi/oα-dependent cAMP reduction and/or Gβγ-dependent modulation of ion channels. In order to analyze some physiological functions, the Ga- and Gβγ-dependent responses need to be separated. For example, opioids induce analgesia as well as various side-effects linked to drug abuse, and these opioid effects depend on signaling pathways driven by Gi/o-coupled opioid receptor(s). Analgesia is mainly caused by Gβγ-dependent GIRK activation, and opioid withdrawal is linked to Gi/oα-dependent cAMP signaling pathway^44,45^. Also, arrestin-dependent signaling pathways are responsible for various side-effects of opioid related to the drug abuse^45,46^. G protein-biased agonists for opioid receptors have been developed in this context, and thus Gβγ-biased drug is desirable but not yet available. Thus, c-opsin1-based optical control tools have the potential to separate Gi/oα- and Gβγ-induced opioid effects. Due to limited spatio-temporal resolution, selective activation of Gβγ-dependent pathways cannot be accomplished by chemogenetic tools as well as G protein-biased ligands for GPCRs, in contrast to c-opsin1. GIRK channels are targets of optogenetics not only in neural cells^7,8^ but also in non-neural tissues such as cardiac atrial cells^40^, and they are linked to various diseases such as Down syndrome, Parkinson’s disease, Keppen–Lubinsky syndrome, and epilepsy^47,48^. Thus, c-opsin1-based optical control tools biased for Gβγ-dependent GIRK channel activation would be useful to understand a wide variety of cellular functions and disease mechanisms.

## Abbreviations

Gi/o: Gi/o-type trimeric G proteins;
GIRK: G-protein-coupled inwardly rectifying potassium channel;
GPCR: G protein-coupled receptor;
LamPP: lamprey parapinopsin;
λmax: absorption maximum;
mouse SWO: mouse short wavelength-sensitive opsin (mouse UV-sensitive cone visual pigment);
UV: ultra-violet;
WT: wild-type

---

**Supplemental Fig. S1.**
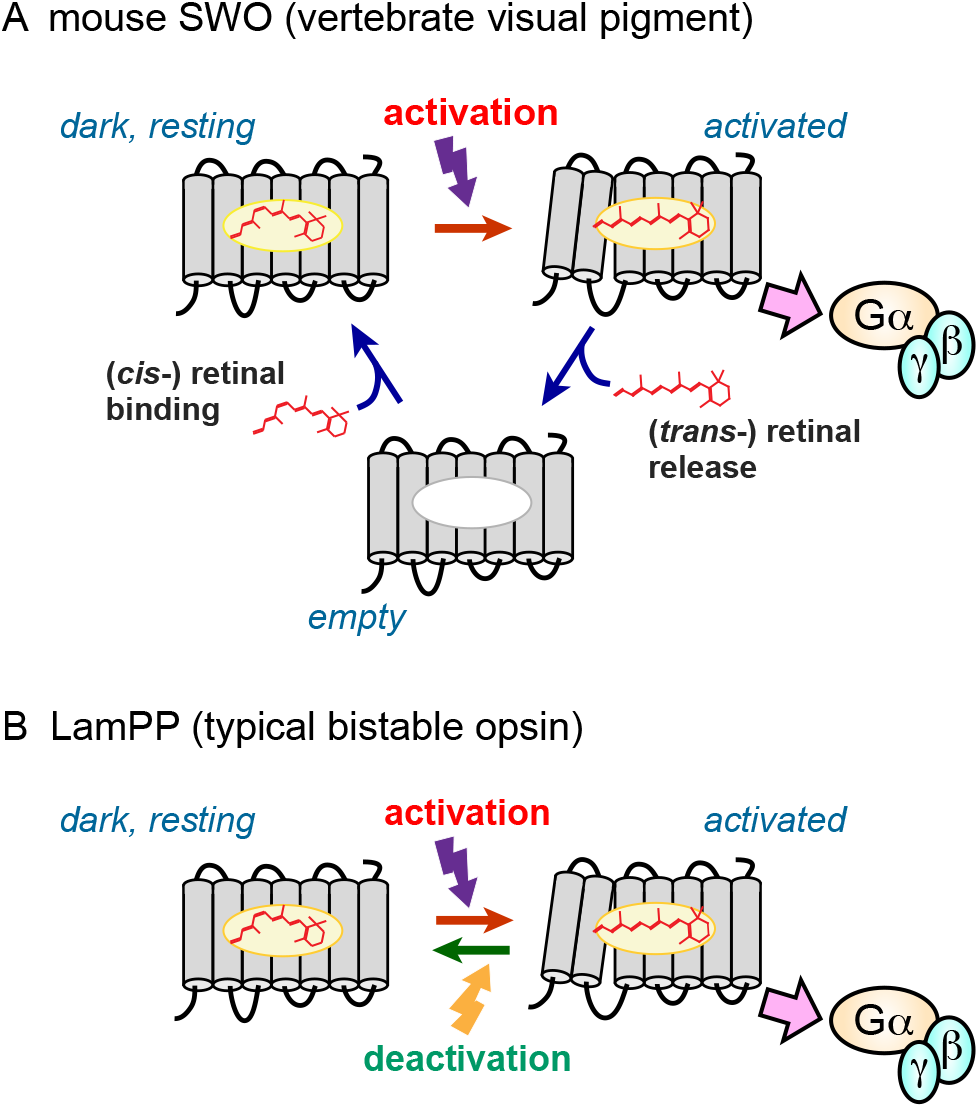
Models describing activation, inactivation, and deactivation processes of opsins used in this study. Schematic drawings of activation/inactivation/deactivation of mouse SWO (vertebrate visual pigments) (*A*), and LamPP (typical bistable opsins) (*B*) are shown. See main text for details.

**<u>Supplemental Data</u> Amino acid sequences of opsins and G proteins used in this study**

**<u>Supplemental Table</u> Summary of molecular properties of opsin constructs as optical control tools**

## Methods

### Ethics statement

All animal experiments in this study were approved by the Animal Care Committee of the National Institutes of Natural Sciences (an umbrella institution of National Institute for Physiological Sciences, Japan), and were performed in accordance with its guidelines.

### Constructs

The cDNAs of *Platynereis* c-opsin1 and the K94T mutant with the coding sequence of the 1D4 tag (ETSQVAPA) on the C-terminus were inserted into the EcoRI/NotI site in a mammalian expression vector pMT and into the EcoRI/HindIII site in the pGEMHE for expression in *Xenopus* oocyte^22^. The plasmids of lamprey parapinopsin (LamPP) with the 1D4 tag in pCDNA3.1^26^, mouse UV-sensitive cone pigment (SWO) with the 1D4 tag in pCDNA3.1^49^ were kindly obtained from Drs. Akihisa Terakita and Mitsumasa Koyanagi (Osaka Metropolitan University), and Dr. Takahiro Yamashita (Kyoto University), respectively, and these cDNA were subcloned into the pMT and pGEMHE vectors. The plasmid of rat RGS8 in pGEMHE^25^ were obtained from Dr. Osamu Saitoh (Nagahama Institute of Bio-Science and Technology), and the cDNA was fused with the sequence of *Platynereis* c-opsin1, a linker sequence (GGGSSGGG), and the 1D4 tag was added on the C-terminus (see Supplemental Data). The cDNA of c-opsin1/RGS8 fusion protein was inserted into the pMT and pGEMHE vectors. The cDNAs of Lg-BiT inserted mouse Giα_2_, human Gβ_1_, and the Sm-BiT fused human Gγ_2_ (C68S) were constructed according to Inoue et al.,^30^ and inserted into the pMT vector. The final amino acid sequences of the constructs are shown in Supplemental Data.

### Preparation of Xenopus oocytes and cRNA injection

*Xenopus* oocytes were isolated from frogs as described previously^22,24,50^. Briefly, *Xenopus* oocytes were surgically collected from frogs anesthetized in water containing 0.15% tricaine and treated with 2 mg/ml collagenase (Sigma-Aldrich) for 3 – 4 h to remove the follicular membrane. 5’-capped cRNA was prepared from the pGEMHE vector containing cDNA of opsin (see above) using an in vitro transcription kit (mMESSAGE mMACHINE Kit, Life Technologies). Typically, we injected the opsin cRNA (~500 pg/oocyte, but for mouse SWO, ~1.5 ng /oocyte) with rat GIRK1 and mouse GIRK2 cRNAs (~25 ng/oocyte and ~12.5 ng/oocyte, respectively) in 50 nL of water/oocyte. The oocytes injected with cRNA were incubated in the standard frog Ringer solution (MBSH), a standard frog Ringer solution (88 mM NaCl, 1 mM KCl, 0.3 mM Ca(NO_3_)2, 0.41 mM CaCl_2_, 0.82 mM MgSO_4_, 2.4 mM NaHCO_3_, and 15 mM HEPES (pH 7.6) with 0.1% penicillin-streptomycin solution), at 17 °C in the dark chamber for 2 days.

### Electrophysiology

We used a conventional two-electrode voltage clamp technique^51,52^ to measure photoresponses caused by opsins. Before electrophysiological recording, oocytes injected with cRNAs were incubated in the standard frog Ringer solution (MBSH) containing 1 μM 11-*cis*-retinal (1/4000 volume of 4 mM retinal in ethanol was added to MBSH) for ~1 h at 17 °C in the dark chamber to form photosensitive pigments. In Figs. 4D, 4F and 5H, 1/4000 volume of ethanol solution instead of retinal solution was added to MBSH. All electrophysiological recordings were performed in a dark room, using only a dim red light which aids handling of oocytes and insertions of electrodes to the oocytes. The red light does not evoke any responses of opsins used in this study. Light-induced electrophysiological responses were recorded in a bath solution (96 mM KCl, 3 mM MgCl_2_, and 5 mM HEPES (pH 7.4)). The tip resistance of the glass electrodes was 0.2–0.5 MΩ when filled with the pipette solution (3 M potassium acetate and 10 mM KCl). The increase in inward K^+^ current as a result of Gi/o activation by opsins was monitored by two-electrode voltage clamp technique using an OC-725C amplifier (Warner Instruments, Hamden, CT) at room temperature with continuous hyperpolarizing pulses of 0.2 s to −100 mV every 2 s from the holding potential of 0 mV and subsequent 0.2-s pluses of 40 mV. In Fig. 5B, hyperpolarizing pulses of 0.2 s to −100 mV every 1 s, because yellow-light induced decrease of GIRK current was very fast and higher time resolution is required to precisely calculate τ_off(+L)_ value. Typically, after 15 repetitions of the hyperpolarizing pulses (~30 s), opsins were activated by illumination with UV light (395-nm LED, Sarspec, Portugal) (light intensity, ~0.7 mW/cm^2^) or blue light (470-nm LED, Sarspec) (light intensity, ~0.3 mW/cm^2^), and subsequently the opsin was deactivated by yellow (>480-nm) light (light intensity, ~30 mW/cm^2^; light source, 3-W LED light, OptoCode, Japan). Data acquisition was performed by a digital converter (Digidata 1440, Molecular Devices, Sunnyvale, CA) and pCLAMP 10 software (Molecular Devices).

### Transfection to COS-1 cells for Glosensor assay and NanoBiT G protein dissociation assay

Opsins, Glosensor assay sensor (coded by pGlo-22F), and NanoBiT-tagged G proteins were transiently expressed in COS-1 cells using polyethyleneimine as described previously^22,53^. For Glosensor assay, each well of 96-well assay plate (Corning, Kennebunk, ME) was transfected with 50 ng opsin plasmid, 50 ng pGlo-22F plasmid (Promega, Madison, WI), and 500 ng polyethyleneimine in 25 μL Opti-mem (Gibco, Waltham, MA) and 75 μL D-MEM (Wako, Japan) containing 10 % (v/v) FBS, 100 units/mL penicillin, and 100 μg/mL streptomycin. For NanoBiT G protein dissociation assay, each well of 96-well assay plate (Corning) was transfected with 50 ng opsin plasmid, 25 ng Lg-BiT inserted Gia2 plasmid, 125 ng Gβ_1_ plasmid, 125 ng Sm-BiT fused Gγ_2_ plasmid, and 500 ng polyethyleneimine in 25 μL Opti-mem (Gibco) and 75 μL D-MEM (Wako) containing 10 % (vol/vol) FBS, 100 units/mL penicillin, and 100 μg/mL streptomycin.

### Glosensor assay

The transfected COS-1 cells were incubated at 37 °C, 5 % CO_2_ for 2 days, and the medium was aspirated, followed by addition of HBSS (145 mM NaCl, 10 mM D-glucose, 5 mM KCl, 1 mM MgCl_2_, 1.7 mM CaCl_2_, 1.5 mM NaHCO_3_, 10 mM HEPES, pH 7.4)^54^ containing 1 μM 11-*cis*-retinal, 2 % (vol/vol) Glosensor cAMP reagent stock solution, and 1 μM forskolin. In Fig. 4, I and J, the final exogenous retinal concentrations were changed to 100 nM or 10 nM. Then, the cells were incubated at room temperature for ~2 h to stabilize luminescence levels. Luminescence was measured using GM-2000 or GM-3510 microplate reader (Promega). Luminescence level of each well was measured every 30 sec with integration time of 0.9 sec. To illuminate opsins, luminescence measurement was interrupted, the plate was ejected, and the plate was illuminated by UV or blue light. UV and blue light source are 375 nm LED (OpotoCode) (light intensity, ~0.5 mW/cm^2^) and 500-nm light (Opto-spectrum generator L12194, Hamamatsu, Japan) (light intensity, ~0.5 mW/cm^2^), respectively. After illumination, luminescence measurement was resumed. The measured luminescence levels were normalized to the level at the starting point (time = 0 min).

### The NanoBiT G protein dissociation assay

The transfected COS-1 cells were incubated at 37 °C, 5 % CO_2_ for 1 day, and the medium was aspirated, followed by addition of HBSS containing 1 μM 11-*cis*-retinal and 5 μM coelenterazine h (Wako). Then, the cells were incubated at room temperature for ~2 h to stabilize luminescence levels. Luminescence was measured using GM-2000 or GM-3510 microplate reader (Promega). Luminescence level of each well was measured every 6 sec with integration time of 0.3 sec. To illuminate opsins, luminescence measurement was interrupted, the plate was ejected, and the plate was illuminated by UV light (375-nm LED, OptoCode) (light intensity, ~0.5 mW/cm^2^). After illumination, luminescence measurement was resumed. The measured luminescence levels were normalized to the level at the starting point (time = 0 min).

## Acknowledgment

We thank Drs. Takushi Shimomura (National Institute for Physiological Science) and I-Shan Chen (National Institute for Physiological Science, Wakayama Medical University) for kind introduction and help of electrophysiological experiments; Drs. Akihisa Terakita, Mitsumasa Koyanagi (Osaka Metropolitan University), Osamu Saitoh (Nagahama Institute of Bio-Science and Technology), and Takahiro Yamashita (Kyoto University) for providing us with plasmids; Dr. David Farrens (Oregon Health & Science University) for providing us with COS-1 cell line and the pMT expression vector. We also thank Chizue Naito (National Institute for Physiological Science), Hiroe Motomura, Kayo Inaba (Institute for Molecular Science), and the Functional Genomics Facility, NIBB Core Research Facilities (Okazaki, Japan) for technical support.

## Author contribution

H. T. designed the study, conducted experiments, and analyzed obtained data. Y. K. supervised electrophysiological experiments. H. T. and Y. K. discussed obtained data and wrote the paper.

---

<u>Supplemental Table</u> Summary of molecular properties of opsin constructs as optical control tools

| opsin construct | light-independent<br>inactivation | light-dependent<br>deactivation | repetition of light-dependent activation<br>and deactivation | Gβγ biased<br>signaling | resistance to<br>retinal depletion |
| --- | --- | --- | --- | --- | --- |
| c-opsin1 WT | ++ | + | + * | + | + |
| LamPP | - | + | + ** | - | N. D. |
| c-opsin1 K94T | + | - *** | - *** | - | N. D. |
| mouse SWO | +++ | - | - | + | - |
| c-opsin1/RGS8 fusion | +++ | ++ | + | ++ | + |

N. D.: not determined in this study
\* Repetitive activation and deactivation of c-opsin1 WT was reported in our previous study (ref. 22).
\*\* Repetitive activation and deactivation of LamPP was reported also in previous studies (refs. 9, 13, 15, 27, and 28).
\*\*\* Due to overlap of absorption spectra of inactive (resting) and activated forms (ref. 22)

## <u>Supplemental Data</u> Amino acid sequences of opsins and G proteins used in this study.

*Platynereis* c-opsin1 with the 1D4 tag

MDGENLTIPNPVTELMDTPINSTYFQNLNAETDGGNHYIYNAFTATDYNICAAYLFFIACLGVSLNVLVLVLFIKDRKLR SPNNFLYVSLALGDLLVAVFGTAFKFIITARKTLLREEDGFCKWYGFITYLGGLAALMTLSVIAFVRCLAVLRLGSFTGLT TRMGVAAMAFIWIYSLAFTLAPLLGWNHYIPEGLATWCSIDWLSDETSDKSYVFAIFIFCFLVPVLIIVVSYGLIYDKVRK VAKTGGSVAKAEREVLRMTLLMVSLFMLAWSPYAVICMLASFGPKDLLHPVATVIPAMFAKSSTMYNPLIYVFMNKQF RRSLKVLLGMGVEDLNSESERATGGTATNQVAATETSQVAPA

Red: the 1D4 tag

*Platynereis* c-opsin1 K94T mutant with the 1D4 tag

MDGENLTIPNPVTELMDTPINSTYFQNLNAETDGGNHYIYNAFTATDYNICAAYLFFIACLGVSLNVLVLVLFIKDRKLR SPNNFLYVSLALGDLLVAVFGTAFTFIITARKTLLREEDGFCKWYGFITYLGGLAALMTLSVIAFVRCLAVLRLGSFTGLT TRMGVAAMAFIWIYSLAFTLAPLLGWNHYIPEGLATWCSIDWLSDETSDKSYVFAIFIFCFLVPVLIIVVSYGLIYDKVRK VAKTGGSVAKAEREVLRMTLLMVSLFMLAWSPYAVICMLASFGPKDLLHPVATVIPAMFAKSSTMYNPLIYVFMNKQF RRSLKVLLGMGVEDLNSESERATGGTATNQVAATETSQVAPA

Red: the 1D4 tag, blue: K94T substitution

*Platynereis* c-opsin1/RGS8 fusion

MDGENLTIPNPVTELMDTPINSTYFQNLNAETDGGNHYIYNAFTATDYNICAAYLFFIACLGVSLNVLVLVLFIKDRKLR SPNNFLYVSLALGDLLVAVFGTAFKFIITARKTLLREEDGFCKWYGFITYLGGLAALMTLSVIAFVRCLAVLRLGSFTGLT TRMGVAAMAFIWIYSLAFTLAPLLGWNHYIPEGLATWCSIDWLSDETSDKSYVFAIFIFCFLVPVLIIVVSYGLIYDKVRK VAKTGGSVAKAEREVLRMTLLMVSLFMLAWSPYAVICMLASFGPKDLLHPVATVIPAMFAKSSTMYNPLIYVFMNKQF RRSLKVLLGMGVEDLNSESERATGGTATNQVAATGGGSSGGGAALLMPRRNKGMRTRLGCLSHKSDSCSDFTAILPDK PNRALKRLSTEEATRWADSFDVLLSHKYGVAAFRAFLKTEFSEENLEFWLACEEFKKTRSTAKLVTKAHRIFEEFVDVQ APREVNIDFQTREATRKNMQEPSLTCFDQAQGKVHSLMEKDSYPRFLRSKMYLDLLSQSQRRLSETSQVAPA

Red: the 1D4 tag, blue: linker, green: RGS8

LamPP with the 1D4 tag

MENLTSLDLLPNGEVPLMPRYGFTILAVIMAVFTIASLVLNSTVVIVTLRHRQLRHPLNFSLVNLAVADLGVTVFGASLVV ETNAVGYFNLGRVGCVIEGFAVAFFGIAALCTIAVIAVDRFVVVCKPLGTLMFTRRHALLGIAWAWLWSFVWNTPPLFG WGSYELEGVRTSCAPDWYSRDPANVSYITSYFAFCFAIPFLVIVVAYGRLMWTLHQVAKLGMGESGSTAKAEAQVSRM VVVMVVAFLVCWLPYALFAMIVVTKPDVYIDPVIATLPMYLTKTSTVYNPIIYIFMNRQFRDCAVPFLLCGRNPWAEPSS ESATAASTSATSVTLASAPGQVSPSETSQVAPA

Red: the 1D4 tag

Mouse SWO with the 1D4 tag

MSGEDDFYLFQNISSVGPWDGPQYHLAPVWAFRLQAAFMGFVFFVGTPLNAIVLVATLHYKKLRQPLNYILVNVSLGG FLFCIFSVFTVFIASCHGYFLFGRHVCALEAFLGSVAGLVTGWSLAFLAFERYVVICKPFGSIRFNSKHALMVVLATWIIG IGVSIPPFFGWSRFIPEGLQCSCGPDWYTVGTKYRSEYYTWFLFIFCFIIPLSLICFSYSQLLRTLRAVAAQQQESATTQKAEREVSHMVVVMVGSFCLCYVPYAALAMYMVNNRNHGLDLRLVTIPAFFSKSSCVYNPIIYCFMNKQFRACILEMVCRKPMADESDVSGSQKTEVSTVSSSKVGPHETSQVAPA

Red: the 1D4 tag

Lg-BiT-inserted mouse Giα_2_

MGCTVSAEDKAAAERSKMIDKNLREDGEKAAREVKLLLLGAGESGKSTIVKQMKIIHEDGYSEEECRQYRAVVYSNTIQSIMAIVKAMGNLGGSGGGGSGGSSSGGVFTLEDFVGDWEQTAAYNLDQVLEQGGVSSLLQNLAVSVTPIQRIVRSGENALKIDIHVIIPYEGLSADQMAQIEEVFKVVYPVDDHHFKVILPYGTLVIDGVTPNMLNYFGRPYEGIAVFDGKKITVTGTLWNGNKIIDERLITPDGSMLFRVTINSGGSGGGGSGGSSSGGQIDFADPQRADDARQLFALSCAAEEQGMLPEDLSGVIRRLWADHGVQACFGRSREYQLNDSAAYYLNDLERIAQSDYIPTQQDVLRTRVKTTGIVETHFTFKDLHFKMFDVGGQRSERKKWIHCFEGVTAIIFCVALSAYDLVLAEDEEMNRMHESMKLFDSICNNKWFTDTSIILFLNKKDLFEEKITQSSLTICF PEYTGANKYDEAASYIQSKFEDLNKRKDTKEIYTHFTCATDTKNVQFVFDAVTDVIIKNNLKDCGLF

Red: Lg-BiT, blue: linker

Human Gβ_1_

MSELDQLRQEAEQLKNQIRDARKACADATLSQITNNIDPVGRIQMRTRRTLRGHLAKIYAMHWGTDSRLLVSASQDGK LIIWDSYTTNKVHAIPLRSSWVMTCAYAPSGNYVACGGLDNICSIYNLKTREGNVRVSRELAGHTGYLSCCRFLDDNQI VTSSGDTTCALWDIETGQQTTTFTGHTGDVMSLSLAPDTRLFVSGACDASAKLWDVREGMCRQTFTGHESDINAICFFP NGNAFATGSDDATCRLFDLRADQELMTYSHDNIICGITSVSFSKSGRLLLAGYDDFNCNVWDALKADRAGVLAGHDN RVSCLGVTDDGMAVATGSWDSFLKIWN

Sm-BiT-fused human Gg2 (with C68S substitution)

MVTGYRLFEEILGGSGGGGSGGSSSGGASNNTASIAQARKLVEQLKMEANIDRIKVSKAAADLMAYCEAHAKEDPLLT PVPASENPFREKKFFSAIL

Red: Sm-BiT, blue: linker, green: C68S substitution

## References

1 Deisseroth, K. Optogenetics. Nat Methods 8, 26–29 (2011).

2 Airan, R. D., Thompson, K. R., Fenno, L. E., Bernstein, H. & Deisseroth, K. Temporally precise in vivo control of intracellular signalling. Nature 458, 1025–1029 (2009).

3 Tsukamoto, H. & Furutani, Y. Optogenetic Modulation of Ion Channels by Photoreceptive Proteins. Adv Exp Med Biol 1293, 73–88 (2021).

4 Spangler, S. M. & Bruchas, M. R. Optogenetic approaches for dissecting neuromodulation and GPCR signaling in neural circuits. Curr Opin Pharmacol 32, 56–70 (2017).

5 Li, X. et al. Fast noninvasive activation and inhibition of neural and network activity by vertebrate rhodopsin and green algae channelrhodopsin. Proc Natl Acad Sci U S A 102, 17816–17821 (2005).

6 Masseck, O. A. et al. Vertebrate cone opsins enable sustained and highly sensitive rapid control of Gi/o signaling in anxiety circuitry. Neuron 81, 1263–1273 (2014).

7 Wiegert, J. S., Mahn, M., Prigge, M., Printz, Y. & Yizhar, O. Silencing Neurons: Tools, Applications, and Experimental Constraints. Neuron 95, 504–529 (2017).

8 Rost, B. R., Wietek, J., Yizhar, O. & Schmitz, D. Optogenetics at the presynapse. Nat Neurosci 25, 984–998 (2022).

9 Koyanagi, M. et al. High-performance optical control of GPCR signaling by bistable animal opsins MosOpn3 and LamPP in a molecular property-dependent manner. Proc Natl Acad Sci U S A 119, e2204341119 (2022).

10 van Wyk, M., Pielecka-Fortuna, J., Lowel, S. & Kleinlogel, S. Restoring the ON Switch in Blind Retinas: Opto-mGluR6, a Next-Generation, Cell-Tailored Optogenetic Tool. PLoS Biol 13, e1002143 (2015).

11 Shichida, Y. & Imai, H. Visual pigment: G-protein-coupled receptor for light signals. Cell Mol Life Sci 54, 1299–1315 (1998).

12 Terakita, A. in Photobiology: Principles, Applications and Effects (eds L. N. Collignon & C. B. Normand) 179–193 (Nova Science Publishers, Inc., 2010).

13 Copits, B. A. et al. A photoswitchable GPCR-based opsin for presynaptic inhibition. Neuron 109, 1791–1809 e1711 (2021).

14 Mahn, M. et al. Efficient optogenetic silencing of neurotransmitter release with a mosquito rhodopsin. Neuron 109, 1621–1635 e1628 (2021).

15 Rodgers, J. et al. Using a bistable animal opsin for switchable and scalable optogenetic inhibition of neurons. EMBO Rep 22, e51866 (2021).

16 Tsukamoto, H. & Terakita, A. Diversity and functional properties of bistable pigments. Photochem Photobiol Sci 9, 1435–1443 (2010).

17 Vierock, J. et al. WiChR, a highly potassium selective channelrhodopsin for low-light one- and two-photon inhibition of excitable cells. Sci Adv, eadd7729 (2022).

18 Nargeot, J. et al. A photoisomerizable muscarinic antagonist. Studies of binding and of conductance relaxations in frog heart. J Gen Physiol 79, 657–678 (1982).

19 Hille, B. Ion Channels of Excitable Membranes. 3rd edn, (Sinauer Associates, 2001).

20 Yatani, A. & Brown, A. M. Rapid beta-adrenergic modulation of cardiac calcium channel currents by a fast G protein pathway. Science 245, 71–74 (1989).

21 Arendt, D., Tessmar-Raible, K., Snyman, H., Dorresteijn, A. W. & Wittbrodt, J. Ciliary photoreceptors with a vertebrate-type opsin in an invertebrate brain. Science 306, 869–871 (2004).

22 Tsukamoto, H., Chen, I. S., Kubo, Y. & Furutani, Y. A ciliary opsin in the brain of a marine annelid zooplankton is ultraviolet-sensitive, and the sensitivity is tuned by a single amino acid residue. J Biol Chem 292, 12971–12980 (2017).

23 Wang, T., Li, Z., Cvijic, M. E., Zhang, L. & Sum, C. S. in Assay Guidance Manual (eds S. Markossian et al. (2004).

24 Kubo, Y., Miyashita, T. & Murata, Y. Structural basis for a Ca2+-sensing function of the metabotropic glutamate receptors. Science 279, 1722–1725 (1998).

25 Saitoh, O., Kubo, Y., Miyatani, Y., Asano, T. & Nakata, H. RGS8 accelerates G-protein-mediated modulation of K+ currents. Nature 390, 525–529 (1997).

26 Koyanagi, M. et al. Bistable UV pigment in the lamprey pineal. Proc Natl Acad Sci U S A 101, 6687–6691 (2004).

27 Eickelbeck, D. et al. Lamprey Parapinopsin (“UVLamP”): a bistable UV-sensitive optogenetic switch for ultrafast control of GPCR pathways. Chembiochem (2019).

28 Kawano-Yamashita, E. et al. Activation of Transducin by Bistable Pigment Parapinopsin in the Pineal Organ of Lower Vertebrates. PLoS One 10, e0141280 (2015).

29 Veedin Rajan, V. B. et al. Seasonal variation in UVA light drives hormonal and behavioural changes in a marine annelid via a ciliary opsin. Nat Ecol Evol 5, 204–218 (2021).

30 Inoue, A. et al. Illuminating G-Protein-Coupling Selectivity of GPCRs. Cell 177, 1933–1947 e1925 (2019).

31 Kato, H. E. et al. Conformational transitions of a neurotensin receptor 1-Gi1 complex. Nature 572, 80–85 (2019).

32 Chen, M. H., Kuemmel, C., Birge, R. R. & Knox, B. E. Rapid release of retinal from a cone visual pigment following photoactivation. Biochemistry 51, 4117–4125 (2012).

33 Hollinger, S. & Hepler, J. R. Cellular regulation of RGS proteins: modulators and integrators of G protein signaling. Pharmacol Rev 54, 527–559 (2002).

34 Ritter, E., Zimmermann, K., Heck, M., Hofmann, K. P. & Bartl, F. J. Transition of rhodopsin into the active metarhodopsin II state opens a new light-induced pathway linked to Schiff base isomerization. J Biol Chem 279, 48102–48111 (2004).

35 Koyanagi, M., Takada, E., Nagata, T., Tsukamoto, H. & Terakita, A. Homologs of vertebrate Opn3 potentially serve as a light sensor in nonphotoreceptive tissue. Proc Natl AcadSci USA 110, 4998–5003 (2013).

36 Korf, H. W., Foster, R. G., Ekstrom, P. & Schalken, J. J. Opsin-like immunoreaction in the retinae and pineal organs of four mammalian species. Cell Tissue Res 242, 645–648 (1985).

37 Kramm, C. M., de Grip, W. J. & Korf, H. W. Rod-opsin immunoreaction in the pineal organ of the pigmented mouse does not indicate the presence of a functional photopigment. Cell Tissue Res 274, 71–78 (1993).

38 Buhr, E. D., Vemaraju, S., Diaz, N., Lang, R. A. & Van Gelder, R. N. Neuropsin (OPN5) Mediates Local Light-Dependent Induction of Circadian Clock Genes and Circadian Photoentrainment in Exposed Murine Skin. Curr Biol 29, 3478–3487 e3474 (2019).

39 Dawydow, A. et al. Channelrhodopsin-2-XXL, a powerful optogenetic tool for low-light applications. Proc Natl Acad Sci U S A 111, 13972–13977 (2014).

40 Cokic, M., Bruegmann, T., Sasse, P. & Malan, D. Optogenetic Stimulation of Gi Signaling Enables Instantaneous Modulation of Cardiomyocyte Pacemaking. Front Physiol 12, 768495 (2021).

41 Makowka, P. et al. Optogenetic stimulation of Gs-signaling in the heart with high spatiotemporal precision. Nat Commun 10, 1281 (2019).

42 Rodriguez-Rozada, S. et al. Aion is a bistable anion-conducting channelrhodopsin that provides temporally extended and reversible neuronal silencing. Commun Biol 5, 687 (2022).

43 Roth, B. L. DREADDs for Neuroscientists. Neuron 89, 683–694 (2016).

44 Koob, G. F., Sanna, P. P. & Bloom, F. E. Neuroscience of addiction. Neuron 21, 467–476 (1998).

45 Kliewer, A. et al. Phosphorylation-deficient G-protein-biased mu-opioid receptors improve analgesia and diminish tolerance but worsen opioid side effects. Nat Commun 10, 367 (2019).

46 Zuo, Z. The role of opioid receptor internalization and beta-arrestins in the development of opioid tolerance. Anesth Analg 101, 728–734 (2005).

47 Luscher, C. & Slesinger, P. A. Emerging roles for G protein-gated inwardly rectifying potassium (GIRK) channels in health and disease. Nat Rev Neurosci 11, 301–315 (2010).

48 Chen, I. S., Eldstrom, J., Fedida, D. & Kubo, Y. A novel ion conducting route besides the central pore in an inherited mutant of G-protein-gated inwardly rectifying K(+) channel. J Physiol 600, 603–622 (2022).

49 Tsutsui, K., Imai, H. & Shichida, Y. Photoisomerization efficiency in UV-absorbing visual pigments: protein-directed isomerization of an unprotonated retinal Schiff base. Biochemistry 46, 6437–6445 (2007).

50 Fujiwara, Y. & Kubo, Y. Ser165 in the second transmembrane region of the Kir2.1 channel determines its susceptibility to blockade by intracellular Mg2+. J Gen Physiol 120, 677–693 (2002).

51 Methfessel, C. et al. Patch clamp measurements on Xenopus laevis oocytes: currents through endogenous channels and implanted acetylcholine receptor and sodium channels. Pflugers Arch 407, 577–588 (1986).

52 Baumgartner, W., Islas, L. & Sigworth, F. J. Two-microelectrode voltage clamp of Xenopus oocytes: voltage errors and compensation for local current flow. Biophys J 77, 1980–1991 (1999).

53 Tsukamoto, H. et al. Retinal Attachment Instability Is Diversified among Mammalian Melanopsins. J Biol Chem 290, 27176–27187 (2015).

54 Goulding, J., May, L. T. & Hill, S. J. Characterisation of endogenous A2A and A2B receptor-mediated cyclic AMP responses in HEK 293 cells using the GloSensor biosensor: Evidence for an allosteric mechanism of action for the A2B-selective antagonist PSB 603. Biochem Pharmacol 147, 55–66 (2018).

